# CLADES — Contrastive Learning Augmented DifferEntial Splicing with Orthologous Positive Pairs

**DOI:** 10.64898/2026.02.20.707118

**Authors:** Arghamitra Talukder, Nicholas Keung, Itsik Pe’er, David A. Knowles

## Abstract

Alternative splicing (AS) reshapes transcript and protein repertoires across biological, e.g. cellular, contexts. However, learning sequence → content-specific splicing mappings is challenging due to limited labels across tissues and cell types and variability introduced by experimental protocols. We propose a contrastive representation learning pre-training approach grounded in evolutionary conservation. Orthologous exon–intron junction sequences are treated as semantically consistent views of the same regulatory program: evolutionary orthologs are positive pairs, non-homologous junctions are negatives. This discriminative objective aligns embeddings of regulatory equivalents while separating functionally unrelated sequences, inducing invariances to unconstrained sequence and emphasizing conserved motif/RBP and positional signals. We show that this pre-training strategy provides representations that help predict *Δψ*, the change in exon inclusion between conditions, which encodes both direction and magnitude of splicing shifts. Specifically, we finetune a lightweight supervised head on available labels to predict *Δψ*. To make these predictions biologically meaningful, we further introduce an interpretable, splice-motif–aware classification framework grounded in known regulatory signals. On benchmarks spanning tissue- and cell-type differential splicing, the learned representations yield strong *Δψ classification* performance (AUPRC/AUROC for increased/decreased inclusion) and competitive results for *regression* (RMSE, Spearman). These findings indicate that evolution-as-augmentation, instantiated via contrastive learning, is an effective and biologically principled route to context-resolved splicing prediction.

## 1 Introduction

Alternative splicing (AS) is a key eukaryotic mechanism for expanding and diversifying transcript and protein repertoires across biological contexts. RNA sequencing has enabled AS quantification at the level of single exon or junction inclusion events, typically reported as percent spliced-in (*ψ*) (13). However, absolute *ψ* in a single context does not by itself reflect regulatory differences; instead, the change in inclusion between biological states, *Δψ*, directly captures context-dependent regulation (23). *Δψ* captures *direction* and *magnitude* of exon-usage changes—the quantities most relevant to differential splicing analyses and biological interpretation—and aligns naturally with designs that compare cell types, tissues, developmental stages, perturbations, or disease states. (19; 24; 25; 22). Differential inclusion is governed by sequence-encoded regulatory programs involving short motifs, their cognate RNA-binding proteins (RBPs), and position-dependent context (2; 5), which together create complex, nonlinear dependencies that are not easily captured by handcrafted features. Neural networks are well suited to capture these nonlinear dependencies and, in doing so, learn enriched embeddings that encode motif–RBP interactions and other contextual features predictive of *Δψ* (7). Building on this capability, prior systems have used end-to-end architectures to predict splicing usage or splice-site activity for variant interpretation (27; 7; 8; 12). While these models predict average *ψ* with high accuracy, accurately predicting *Δψ* between tissues or cell-types from sequence remains an open challenge.

To capture the distinctive and functionally relevant feature representations of data, contrastive learning (CL) offers a general framework for representation learning (6). CL learns embeddings by bringing together related inputs and pushing apart unrelated ones. This method falls within self-supervised learning, leveraging self-generated targets from the sequences to learn representations that generalize to downstream analyses absent external annotation (20; 9). This self-supervised strategy reduces our dependence on expensive and noisy experimental annotation, and potentially builds a more robust representation that is less prone to overfitting on spurious, protocol-specific artifacts common in functional genomic data, e.g. GC content bias (10).

We introduce **CLADES**(Contrastive Learning Augmented DifferEntial Splicing), a contrastive learning framework designed to model the sequence determinants of splicing regulation grounded in evolutionary conservation—hypothesizing that the regulatory programs governing context-dependent exon inclusion are deeply conserved. This insight shapes the core contributions of our framework:

### Leverage orthologous positive pairs

A discriminative CL framework is particularly well-suited for genomic data, as it allows evolutionary information to be incorporated through orthologous positive pairs (OPP) as a powerful augmentation strategy (18). This is critical because CL relies on data augmentation to create semantically-related “views” of an input, which serve as its positive pairs (11; 6). In computer vision, for instance, these views are often just different crops or color jitters of the same image. We posit that OPP sequences are semantically consistent “views” of the same underlying regulatory program because evolution preserves function, if not sequence. For example, an exon that is important for neuronal function but detrimental if included in a heart cell needs *cis* regulatory sequences that encode this function. This functional information is implicitly transmitted and conserved through evolution: heredity passes sequence through a noisy mutation–drift channel, yet stabilizing selection preserves the regulatory features that encode conserved phenotypes. Sequence representations that maximally agree between orthologous junctions will therefore encode AS function (18; 1). We thus treat orthologous junction sequences as positive pairs (and non-homologous junctions as negatives). We rely on the abundance of aligned genomes across a large number of vertebrate species[ref], that contribute information through their evolutionary relationship to human, despite the scarcity of context-specific RNA-seq data in most of them. While this assumption generally holds for conserved regulatory programs, it may not apply in cases where splicing phenotypes have diverged across lineages; such instances represent exceptions where orthologous relationships no longer imply shared regulatory function as discussed more extensively in Appendix, Section 1.1.

### Enable generalization to downstream tasks without tissue-specific supervision

Because the framework learns directly from evolutionary relationships rather than condition-specific annotations, it captures general regulatory principles of splicing regulation that extend across tissues and cell-types instead of dataset-specific biases. UMAP projections reveal that exon embeddings organize by both average inclusion level and tissue-specific AS pattern, confirming that CLADES learns biologically meaningful structure. This property allows CLADES to transfer its learned representations to diverse downstream prediction tasks with minimum fine-tuning. On the ASCOT dataset spanning 56 human tissues, CLADES consistently outperforms prior state-of-the-art (SOTA) model MTSplice (7) in predicting tissue-dependent splicing changes. Across nearly all tissues, CLADES shows higher Spearman correlation (*ρ*) between predicted and observed *Δψ* values. On the Tabula Sapiens dataset of 112 cell types, CLADES achieves robust performance. In addition to regression, we further reformulate differential splicing as a classification problem by introducing the Tissue-Specific Regulation Classification (TSRC) and Exon-Level Regulation Classification (ELRC) tasks, which frame *Δψ* prediction in terms of regulatory directionality. CLADES yields higher precision, F1-score, AUPRC, and AUROC for both highly and weakly included exons, indicating that its representations capture transferable regulatory signals and generalize across biological contexts without tissue-specific supervision.

## 2 Methods

We introduce the key notations used in this work in Appendix, Table A1 and present an overview of our framework in Figure 1. This section describes the pretraining and fine-tuning datasets, the corresponding model architectures, and the training objectives used at each stage.

**Fig. 1:**
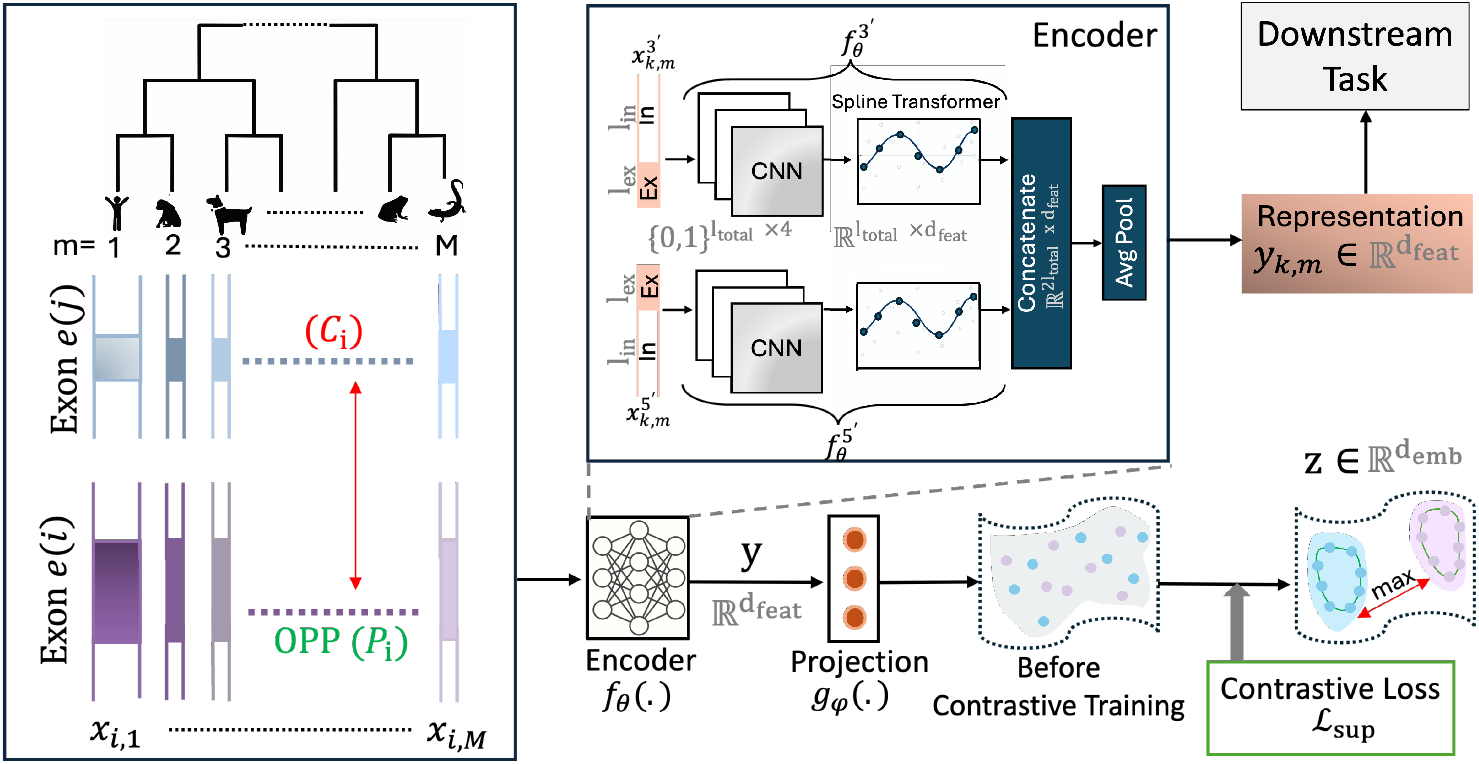
Overview of the framework. Orthologous positive exon pairs (OPPs) across species are encoded via parallel CNN–Spline branches, pooled into exon-level representations, and projected into an embedding space where a supervised contrastive loss aligns orthologous exons while separating unrelated ones. The representations from pretrained encoder is then used for downstream *Δψ* prediction tasks.

### 2.1 Data Processing

The pretraining dataset consists of orthologous exons and their flanking intronic regions across multiple species, used to learn evolutionary and regulatory representations through contrastive learning. For finetuning, we used the ASCOT dataset (16) to predict tissue-specific differential exon inclusion (*Δψ*) and the Tabula Sapiens dataset (21) to extend to cell-type–specific *Δψ*.

#### Pretraining Dataset

The pretraining dataset was constructed from the Multiz100way (3) multiple sequence alignment (MSA) of vertebrate genomes available at (14), (Appendix, Figure A4). We extracted exon–intron boundary coordinates from the UCSC knownGene annotations and used them to identify orthologous exon alignments across species. For each aligned exon, we included only those species with reference gene annotations available in the UCSC *refGene* or *knownGene* FASTA files and retrieved their corresponding nucleotide sequences. Lineage-specific or non-conserved regions naturally correspond to cases lacking cross-species alignment and are therefore not introduced as training signals. Figure 2 shows each species with their corresponding exon numbers. The dataset was randomly divided into training (80%, *n* = 739,325) and validation (10%, *n* = 92,420) exons. Importantly, exons belonging to the ASCOT fine-tuning test hold-out set were explicitly excluded from the pretraining dataset to prevent any overlap between representation learning and downstream evaluation, thereby avoiding data leakage.

**Fig. 2:**
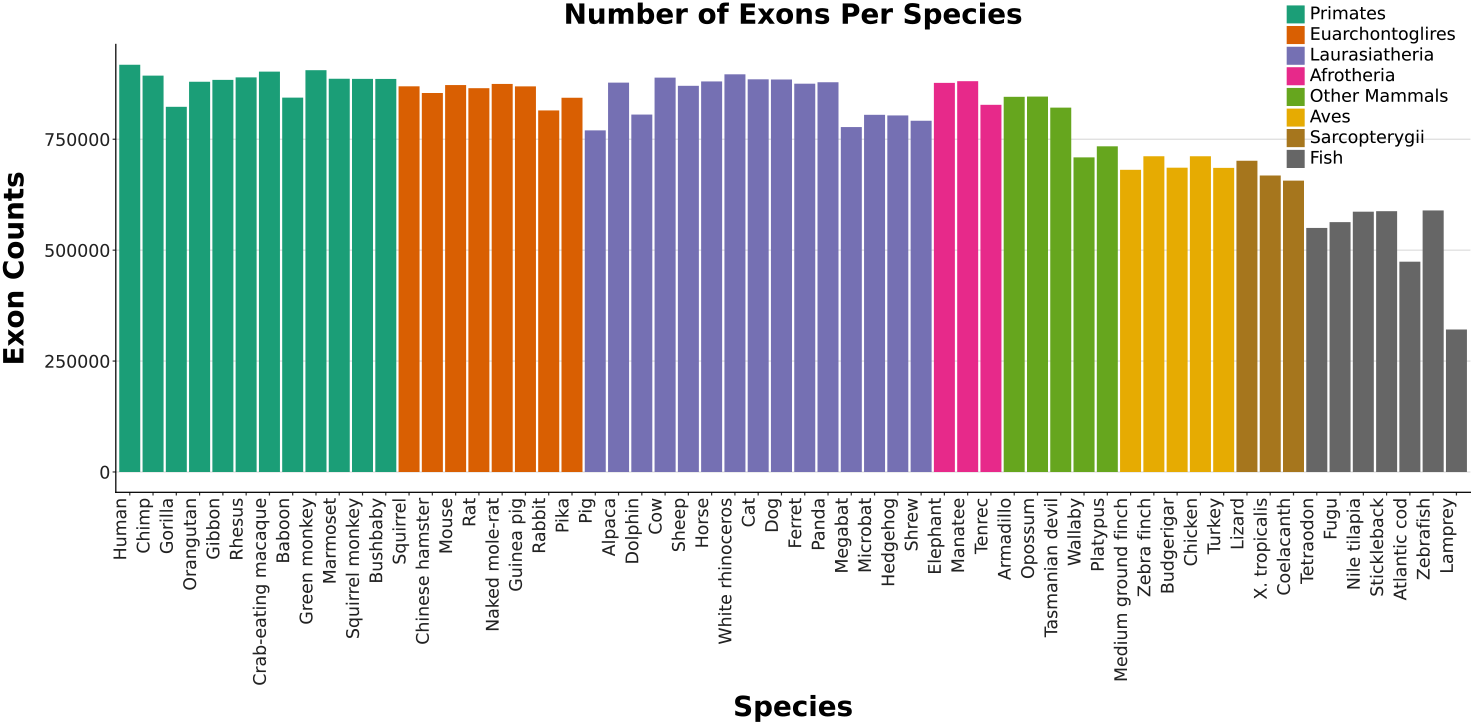
Exon counts across species used for constructing OPPs. Each bar shows the total number of annotated exons used for a given species from the Multiz100way alignment. Species are grouped by major evolutionary clades.

#### Finetuning Dataset

*The ASCOT tissue-specific splicing dataset* was used for fine-tuning, using the same training/validation/test splits as in MTSplice (7). It comprises 61,823 cassette exons with tissue-specific splicing profiles across 56 human tissues from the GTEx dataset (17) (Appendix, Figure A5). The dataset was split by chromosome: 38,028 exons (approximately 61.5%) from chromosomes 4, 6, 8, 10–23, and the sex chromosomes were used for training; and 11,956 exons (approximately 19.3%) from chromosomes 1, 7, and 9 for validation. Testing was performed on 1,621 variable exons drawn from chromosomes 2, 3, and 5, independent held-out set. Variable exons were defined as those whose tissue-specific inclusion (*ψ*_*e,t*_) deviated from their tissue-averaged 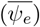 by at least 0.2 in at least one tissue, with the corresponding gene expressed in at least ten tissues.

*The Tabula Sapiens cell-specific dataset* was used as the source of single-cell RNA sequencing data for this study ((21). We were able to obtain 40,884 cells from 17 donors, including cell types that mapped to 199 Cell Ontology classes. Cell types with less than 30 cells were removed, leaving 112 cell types in our dataset (Appendix, Figure A6). The scRNA-seq data were then combined according to cell type, and we identified alternative splicing events for each cell type using rMATS-turbo (26). From the junction counts for skipped exon events, we calculated *ψ*_*e,t*_ for each exon and calculated 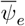 across each cell type. In total, 61,846 exons (54.08%) exhibited at least one cell type with a *ψ* value deviating from 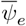 by ≥ 10 percentage points. In addition, 20,145 exons (17.6%) differed from the mean in 10 or more cell types (Appendix, Figure A7, A8). Our final data contained 114,366 total exons across 112 cell types. The data was split randomly, with 80,058 exons in the training set (70%), 17,154 exons in the validation set (15%), and 17,154 (15%) exons in the test set.

### 2.2 Contrastive Learning Pre-training and Finetuning

#### Contrastive Learning Objective

During the pretraining phase, our goal is to learn biologically meaningful representations of exon–intron sequences that capture conserved regulatory features across species. As input, we used *l*_in_ = 300 bp of exon-adjacent intronic sequence and *l*_ex_ = 100 bp of exonic sequence from both the upstream (5*′*) and downstream (3*′*) ends of each exon, as illustrated in Figure 1. For each minibatch, we sample *N* human “anchor” exons, each associated with a set of orthologous exons from other species. Specifically, for each anchor exon *e*_*i*_ (*i* = 1, …, *N*), we sample *M* orthologous exons (OPPs), i.e., 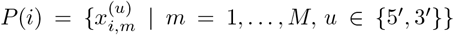, while samples from other anchors {*P* (*j*)}_*j*≠*i*_ serve as negatives in the contrastive objective. Each input sequence *x*_*i,m*_ consists of two fragments corresponding to the upstream (5*′*) and downstream (3*′*) exon–intron boundaries, each represented as a one-hot matrix 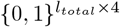 where *l*_*total*_ = *l*_*in*_ + *l*_*ex*_ = 400 The two fragments are encoded independently by 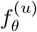 with *u* ∈ {5^*′*^, 3^*′*^}, yielding feature maps 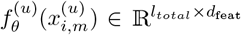, where *d*_feat_ = 64 denotes the feature dimension of each boundary encoder. These representations are concatenated along the sequence dimension to form a joint feature map of size 2*l*_*total*_ *× d*_feat_, which is then globally averaged to obtain a representation 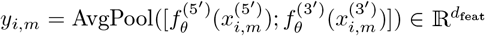. The pooled representation is passed through a two-layer projection head 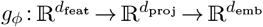, producing the final embedding 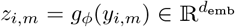, where *d*_proj_ = 512 and *d*_emb_ = 128.

For each anchor exon *e*_*i*_, we define the set of OPP examples *P*(*i*) as all homologous sequences associated with *e*_*i*_. The set of all possible contrastive samples *C*(*i*) is defined as all embeddings within the current batch excluding those of the same anchor, i.e., 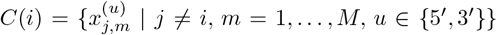. To this end, we employ a supervised contrastive learning objective (15), which generalizes the standard instance discrimination framework to allow multiple positive examples per anchor. The supervised contrastive loss for a batch of *N* anchor exons is,

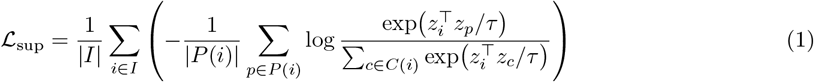

where *P*(*i*) and *C*(*i*) denote the sets of positive and contrastive samples for the *i*-th anchor, and *τ* is the temperature hyperparameter that scales similarity scores. Unlike traditional self-supervised formulations (e.g., NT-Xent loss (6)), which rely on two augmented views of each sample, the supervised contrastive objective allows us to incorporate multiple homologous sequences as augmentations per anchor. After hyperparameter tuning, we selected the configuration with the lowest validation loss: temperature *τ* = 0.2, batch size *N* = 2048 and *M* = 10 augmentations. Training was performed for 25 epochs on EmpireAI (4) nodes equipped with H100 GPUs.

#### Encoder and fine-tuning architecture

The encoder design follows the MTSplice (7) architecture, which employs parallel (for 5’ and 3’ context) convolutional and spline transformation layers to capture position-dependent sequence motifs. For fine-tuning, the encoder output is passed through two fully connected layers with batch normalization and dropout regularization. The final model outputs a real-valued vector *ŷ*_*i*_ ∈ℝ^*T*^ for each exon *e*, representing its tissue- or cell-type–specific logit of inclusion level *Δ*logit(*ψ*_*e,t*_), where *T* denotes the number of biological contexts (tissues or cell types). During training, the predicted differential logits are added to the logit of the mean inclusion 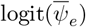 per exon, where the mean value is derived from the ground-truth data. A sigmoid activation *σ*(·) is applied to obtain tissue-specific inclusion probabilities,

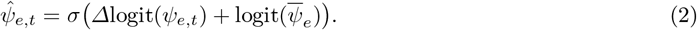

During fine-tuning, the model is trained to minimize the Kullback–Leibler (KL) divergence between the predicted and observed tissue-specific inclusion levels (*ψ*),

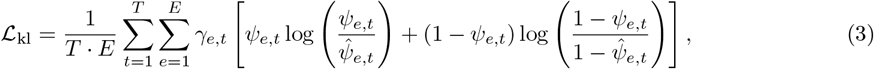

where *γ*_*e,t*_ ∈ {0, 1} indicates whether *ψ*_*e,t*_ is observed.

Twenty models were trained for 5 epochs with identical settings but different initializations. An ensemble was constructed through forward selection based on validation loss, sequentially adding models as long as the loss continued to decrease. The final ensemble prediction was obtained by averaging the selected models.

## 3 Results

We evaluate on two complementary resources—the ASCOT variable-cassette-exon benchmark at the tissue level and the Tabula Sapiens compendium at the cell-type level—using the *Δψ* target. Performance is evaluated using Spearman correlation (*ρ*) to assess rank consistency, RMSE to measure effect magnitude accuracy in logit scale, and classification to detect up-vs. down-regulation. Across both datasets, our context-aware sequence representation achieves strong classification performance for direction-of-change detection, with competitive regression accuracy on *Δψ*, indicating that the learned features capture regulatory signals that are transferable across species, tissues, and cell types.

### Tissue-Specific Evaluation on ASCOT Dataset

We evaluated CLADES on the ASCOT dataset spanning 56 human tissues, comparing its performance against the state-of-the-art (SOTA) MTSplice (7) model to assess how well it predicts tissue-dependent splicing changes. Firstly, we examined whether the learned exon embeddings from our contrastively trained encoder reflect biologically meaningful patterns of inclusion and regulation. Figure 3(a) highlights four representative cassette exons—*TIAM2* (exon 23), *FUNDC1* (exon 2), *ASCC1* (exon 8), and *KCNU1* (exon 25)—along with their orthologous counterparts across species, illustrating how exon-specific homologs cluster distinctly in UMAP space. These groupings indicate that OPPs share similar sequence features, which could be related to regulatory similarity (UMAP-x: *H* = 129.08, *p <* 10^−27^; UMAP-y: *H* = 106.94, *p <* 10^−22^; Kruskal–Wallis test). Next, Figure 3(b) presents a UMAP projection of exon embeddings, colored by their mean inclusion level 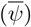. A clear enrichment is observed between exons with high 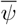 and those with low 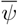, indicating that the embedding space organizes exons with correlation to inclusion propensity; (UMAP-x: *U* = 99,368, *p <* 10^−8^; UMAP-y: *U* = 120,858, *p* = 0.36). Within a tissue group (e.g., brain), exons were stratified by how strongly their *ψ*_*e,t*_ deviated from the global mean: those with elevated inclusion were classified as up-regulated and those with reduced inclusion relative to their average were classified as down-regulated. These groups show enrichment in specific regions of UMAP space Figure 3(c); UMAP-x: *U* = 76,222, *p <* 10^−16^; UMAP-y: *U* = 126,992, *p <* 5 × 10^−4^). Figure 3(d) illustrates how contrastive saliency reveals the specific splice-site features that distinguish high- and low-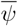 exons. We contrasted embeddings from 20 high- and 20 low-inclusion exons and used the gradient of their negative cosine similarity to highlight discriminative sequence positions (*l*_*in*_ = 200 bp, and *l*_*ex*_ = 100 bp). The final saliency score was obtained by averaging these gradients. This quantity identifies the nucleotides whose perturbation most strongly increases the dissimilarity between the two exon groups, thereby highlighting the positions that drive contrastive separation. The resulting saliency trace exhibits a clear spatial structure: saliency is highest exactly at the splice junctions and decays rapidly as we move farther into the flanking introns, indicating that the model relies predominantly on cis-regulatory signals around the splice junctions. Zooming into these peak regions, motif logos extracted from the upstream and downstream windows reveal strong enrichment of canonical splice-site elements. At the splice acceptor (left zoom), we observe the characteristic **AG** motif, whereas at the splice donor (right zoom), we recover the canonical **GT** dinucleotide followed by purine-rich positions. These motifs confirm that CLADES attends to biologically meaningful splice-site features when distinguishing exons with consistently high versus low inclusion.

**Fig. 3:**
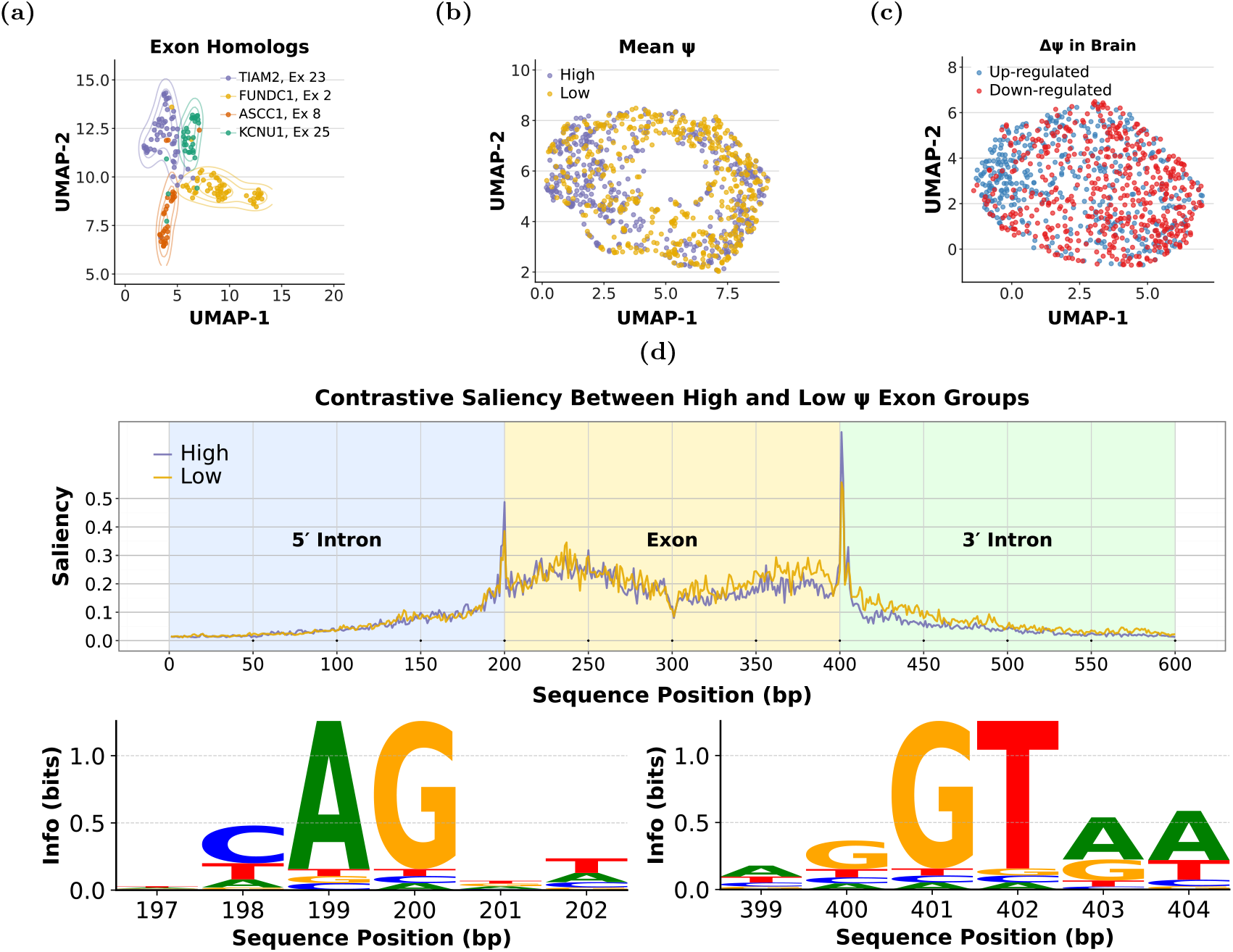
Contrastive embedding structure, tissue-level variability, and saliency-driven motif discovery. **(a)** UMAP projection of exon embeddings learned through contrastive pretraining, showing that homologous exons cluster tightly together across species. **(b)** UMAP of exon embeddings colored by mean *ψ*, distinguishing high- and low-inclusion exons. **(c)** UMAP of *Δψ* in brain, highlighting exons up-regulated (blue) or down-regulated (red) relative to baseline inclusion. **(d)** Contrastive saliency profile across the full exon–intron window for high- and low-*ψ* exon groups. Saliency sharply peaks at the exon boundaries, indicating that the model relies most heavily on the canonical splice-site dinucleotides. Motif logos extracted from these peak positions reveal strong enrichment of the conserved **AG** motif at the upstream splice acceptor and the canonical **GT** dinucleotide at the downstream splice donor.

We therefore next assess whether these learned regulatory features translate into improved tissue-level splicing prediction on the ASCOT dataset. Figure 4(a) shows the *ρ* between predicted and observed *Δψ* values for each tissue. Across nearly all tissues, CLADES outperforms previous SOTA model, indicating that its learned representations generalize better to diverse biological contexts. As an example, Figure 4(b) shows predictions for testis, where the model achieves a strong match with the ground truth (*ρ* = 0.80). To examine how sample availability affects performance, we grouped tissues by the number of exons with valid *ψ* measurements into three categories—low, medium, and high—representing increasing data richness (Appendix, Figure A5). Tissue-category cutoffs were set one standard deviation below and above the mean number of exons. Figure 4(c) summarizes the mean *ρ* within each group. Table 4(d) shows the total number of tissue in each category (**N**_tis_) with the average exon number 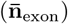 evaluated for that category. CLADES performs consistently better than SOTA in the low group while also matching it in the medium and high groups. CLADES performs especially well in tissues with fewer data points. For well-sampled tissues, both CLADES and SOTA perform similarly, as abundant data already supports tissue-specific learning.

**Fig. 4:**
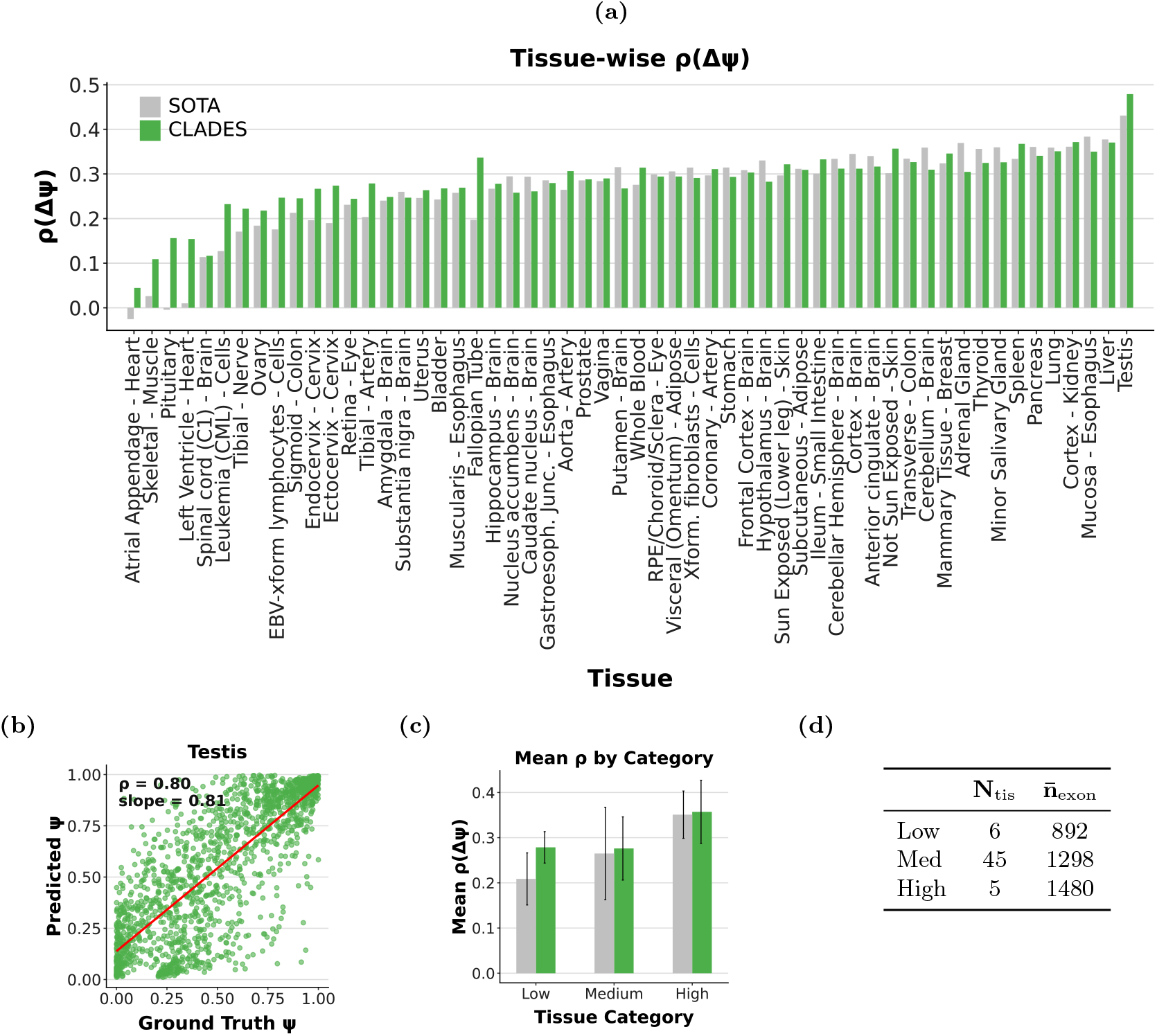
Comparison of CLADES and MTSplice across tissue-specific splicing benchmarks. **(a)** Tissue-wise Spearman correlation (*ρ*) between predicted and observed *Δψ* across all tissues, comparing CLADES with the SOTA model MTSplice. **(b)** Scatter plot for *Testis* between predicted versus ground truth (*ψ*). **(c)** Mean Spearman correlation (*ρ*) of *Δψ* across tissues grouped by sample size (low, medium, high). Error bars indicate ± 1 standard deviation across tissues. **(d)** Summary of tissue-category statistics used in panel (c). For each tissue category, the table reports the number of tissues (**N**_tis_) and the average number of exons 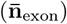 available for evaluation.

Although *Δψ* is a continuous quantity, its biological interpretation is inherently directional: in a given tissue, an exon may be up-regulated, down-regulated, or remain approximately unchanged w.r.t. its default inclusion level 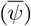. These three regulatory regimes capture the qualitative outcome of tissue-specific splicing, motivating a complementary classification formulation focused on predicting the direction of change. We refer to this as the *Tissue-Specific Regulation Classification (TSRC)* task, which evaluates whether the model can correctly determine the regulatory state of an exon—i.e., whether it is up-regulated, down-regulated, or unchanged—in each tissue relative to its baseline inclusion level. To define these classes systematically, we compared each exon’s tissue-specific *ψ* value to its mean inclusion level across all tissues, computing the differential inclusion as 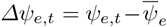. Following a *Tissue-Specific Regulation Labeling (TSRL)* procedure, we labeled each exon **negative** (down-regulated, *Δψ*^−^), **unchanged, (***Δψ*_∅_), or **positive** (up-regulated, *Δψ*^+^) based on its differential inclusion level *Δψ*_*e,t*_, with cutoffs ± *σ*_*e*_ set at one standard deviation away from the mean. This procedure resulted in 11.1% up-regulated, 9.6% down-regulated, and 79.3% unchanged samples.

Table 1 summarizes the model performance across a range of architectural and training configurations. It compares different input configurations (intron-only vs. intron+exon), sequence lengths, and the number of OPPs used during contrastive learning (5 or 10). For intron-only, we used *l*_*in*_ = 200 or 300 bp windows adjacent to the exon, whereas the intron+exon inputs also included *l*_*ex*_ = 100 bp of splice site-adjacent exonic sequence. The table also includes model comparisons trained with and without contrastive learning while keeping the encoder architecture fixed. The SOTA model serves as the non-contrastive baseline for the intron+exon configuration, and we trained an additional intron-only model without contrastive learning pretraining as the baseline for intron-only. Performance is reported in terms of rank consistency (*ρ*), classification accuracy for distinguishing up- and down-regulated exons from unchanged ones (AUPRC/AUROC), and regression error in effect magnitude (RMSE). Across all settings, CLADES achieves the best *ρ* and RMSE with the 200 bp intron+exon input and 10-augmentation configuration. This setting yields a 16% improvement in correlation and a 0.7% reduction in error relative to the SOTA model. CLADES also shows higher *Δψ*^+^ vs *Δψ*_∅_ AUPRC and AUROC values than SOTA, reflecting a clearer distinction between up-regulated and unchanged exons while maintaining comparable performance for down-regulated events. Focusing on the 300 bp intron+exon configuration CLADES achieves strong classification performance for both up- and down-regulated exons. The best (*Δψ*^+^ vs *Δψ*_∅_) represents a 14% increase in AUPRC and a 9.7% increase in AUROC compared to the SOTA model. The 200 bp intron+exon window performs best for regression (Spearman *ρ*, RMSE) because it concentrates the model on highly informative splice-proximal regulatory elements, reducing noise and yielding a smoother, more stable mapping from sequence to *Δψ*. The 300 bp window incorporates additional distal intronic context that, while noisier for fine-grained effect-size prediction, provides extra discriminative signals that help separate strongly up- or down-regulated exons from unchanged ones, leading to higher AUPRC and AUROC for classification. Thus, shorter windows favor quantitative accuracy, whereas longer windows enhance sensitivity to large regulatory shifts. In contrast, the intron-only models achieve comparable classification accuracy but show noticeably lower *ρ*, indicating that while they can identify direction of change, they are less effective at modeling the precise magnitude of splicing differences. Since intron-only inputs already lack exon-derived regulatory information, their representational capacity is inherently limited. However, when trained with contrastive learning, these models still achieve competitive performance, suggesting that the CL objective helps recover meaningful structure from purely intronic context. For example, the 200 bp intron-only model improves its Spearman correlation from *ρ* = 0.188 without CL to *ρ* = 0.268 with CL. Across all input types, increasing the number of augmentations from 5 to 10 slightly improves rank correlation and stabilizes RMSE, suggesting that additional contrastive views provide modest but consistent gains in representation quality. Overall, CLADES’ 300 bp intron+exon model offers the best balance between correlation, classification accuracy across tissues.

**Table 1:**
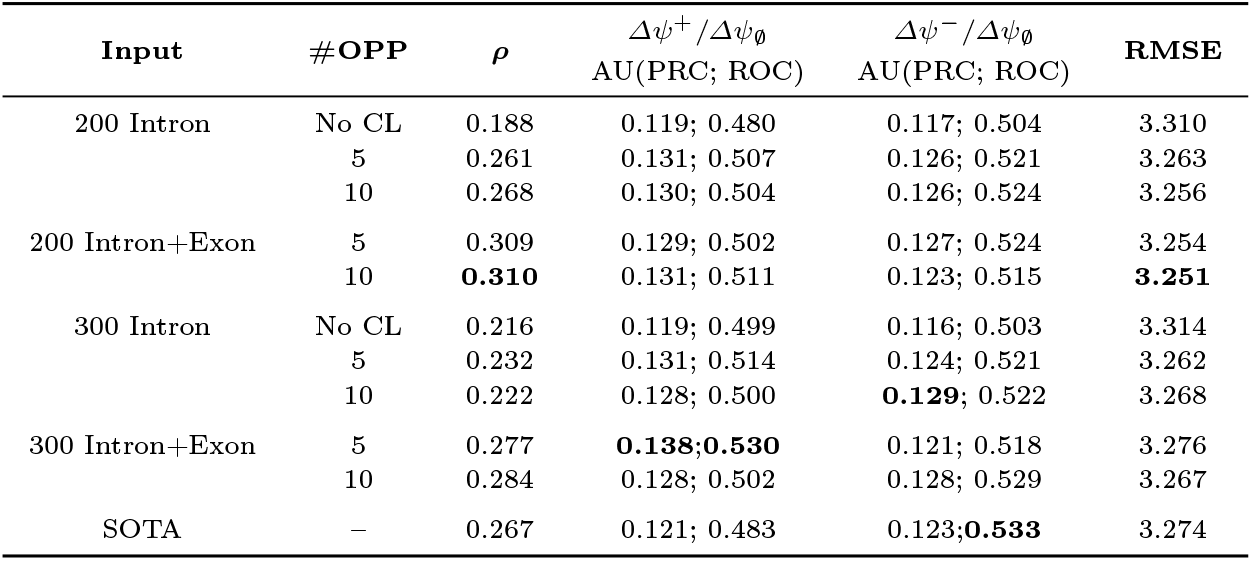
ASCOT TSRC Performance Across Input Configurations and OPP Numbers. Performance comparison on the ASCOT dataset across different input configurations (Intron, Intron+Exon) and OPP strategies, in the Tissue-Specific Regulation Classification (TSRC) task. Reported values include mean performance for *Δψ* predictions: Spearman correlation (*ρ*), precision–recall and ROC scores for up-regulated (*Δψ*^+^) vs. unchanged (*Δψ*_∅_) and down-regulated (*Δψ*^−^) vs. unchanged (*Δψ*_∅_) comparisons, and RMSE. Bold values indicate the best performance within each metric group.

### Cell-Specific Evaluation on Tabula Sapiens Dataset

We next assessed CLADES’ performance at the single-cell-type level using the Tabula Sapiens compendium, which includes 112 distinct cell types spanning multiple tissues. Here, we compared the 300 bp intron+exon, 10-augmentation CLADES model against the same architecture trained without contrastive pretraining as baseline.

Figure 5(a) shows the correlation between predicted and observed *ψ* values for Basal Cells, where CLADES achieves a *ρ* of 0.75. Across all 112 cell types, we computed cross-cell-type *ρ*(*Δψ*) and summarized them by sample size category in Figure 5(b). The cell categorization by sample size follows the same procedure used for the ASCOT tissues in Section 3 (Appendix, Figure A6). CLADES performs slightly below the baseline in cell classes with limited samples, but its advantage becomes clear as the number of available exons increases—showing steadily higher correlations in medium- and high-sample groups. Table 5(c) summarizes numbers of cell types and average exons analyzed within each group 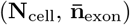.

**Fig. 5:**
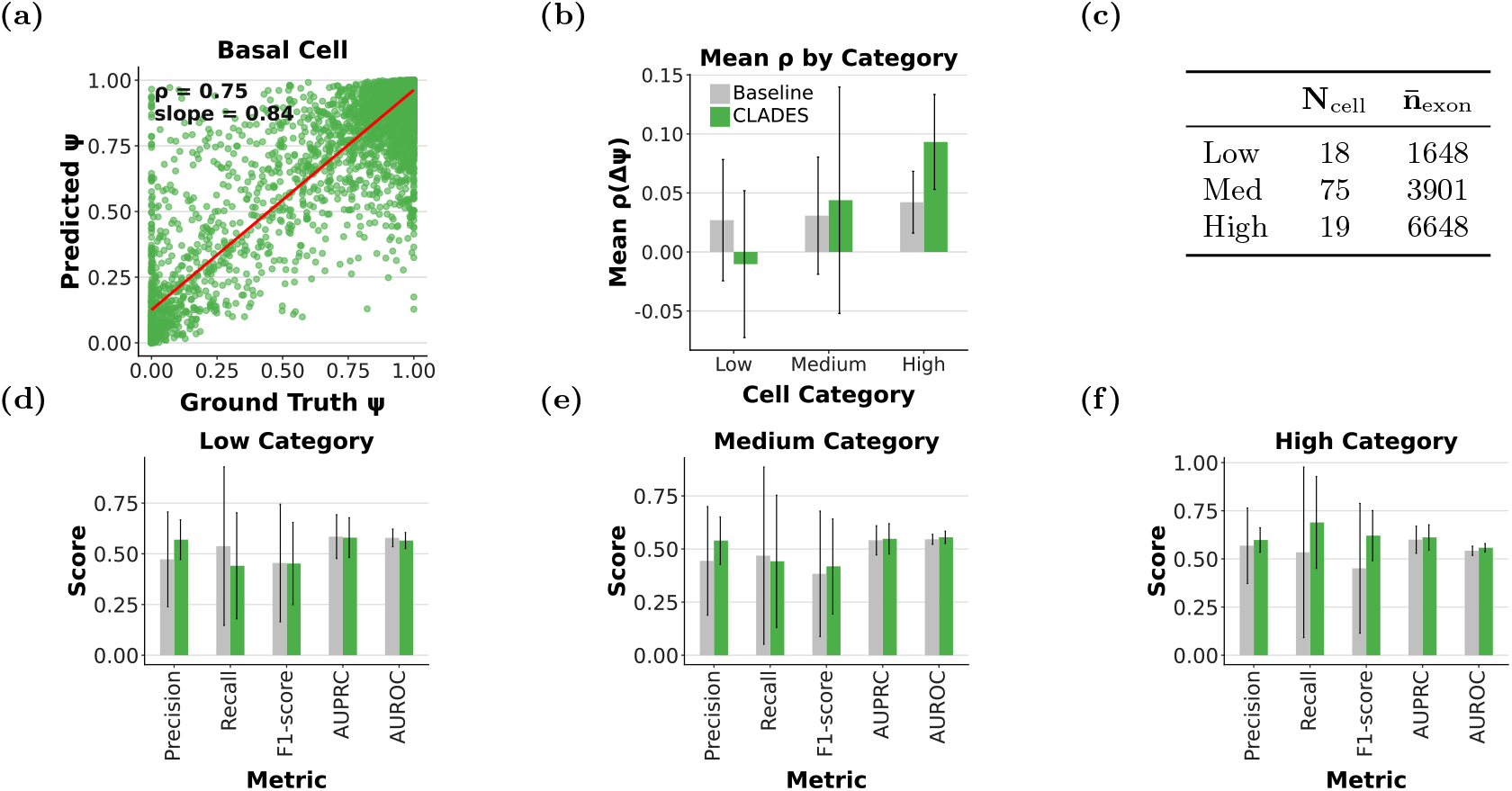
Cell-type–specific evaluation of CLADES on the Tabula Sapiens dataset. Bars showing mean ± standard deviation across cell-type categories. **(a)** Example of predicted versus ground truth *ψ* for the *Basal Cell* type. **(b)** Mean Spearman correlation (*ρ*) of *Δψ* across cell-type categories (Low, Medium, High), comparing CLADES with the baseline model. **(c)** Summary of cell-type categories showing the number of cell types (**N**_cell_) and the mean number of exons 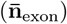 per category. **(d–f)** Classification performance across cell-type categories (Low, Medium, High) based on Precision, Recall, F1-score, AUPRC, and AUROC metrics.

To assess the TSRC task, we followed the same TSRL protocol, with the only modification being the use of a stricter threshold of 1.5 × *σ*_*e*_ to define up- and down-regulated exons. ASCOT evaluates variable exons with substantial across-tissue variability (Section 2.1), whereas Tabula Sapiens includes randomly selected exons with much lower variance. Accordingly, we used a stricter threshold for classification.

This resulted in 94.3% unchanged, 4.61% down-regulated, and 1.09% up-regulated exon–cell pairs. Because single-cell data are inherently noisy and cell-type definitions are more granular, we focused on the simpler task of distinguishing up-(*Δψ*^+^) versus down-regulated (*Δψ*^−^) exons. As shown in Figure 5(d–f), CLADES outperforms the baseline across all groups and performs best in high-sample classes, surpassing the baseline in every metric. To further examine whether these improvements extend to individual exons, we next evaluated CLADES’ performance at comparing how well it detects differential inclusion patterns among highly and weakly expressed exons. We refer to this as the *Exon-Level Regulation Classification (ELRC)* task.

The mean inclusion 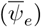 was considered high if it exceeded 0.5 (15,030 exons) and low otherwise (2,126 exons). For high-mean-inclusion exons, our goal is to assess whether the model can detect context-dependent repression, i.e., whether it can recognize when an exon that is usually highly included exhibits a drop in inclusion in a specific cell type. Accordingly, we labeled a high-mean-inclusion exon as positive in a given cell type if its observed inclusion (*ψ*_*e,t*_) was substantially below its baseline, specifically if *ψ*_*e,t*_ *<* 0.6, which resulted in 0.77% of high-mean-inclusion observations being labeled as positive. Conversely, low-mean-inclusion exons were labeled as positive when activated, i.e., if (*ψ*_*e,t*_ *>* 0.4, resulting in 2.7% of them as positives. The remaining exons were labeled neutral (“0”). Table 2 compares classification performance between CLADES and the baseline model across exons with high and low mean inclusion. For highly included exons, CLADES achieved relative improvements of 12.0% (Precision), 6.0% (F1-score), 5.0% (AUPRC), and 4.3% (AUROC) while maintaining comparable Recall. For weakly included exons, CLADES achieved relative improvements of 6.1% (Precision), 6.4% (Recall), 6.5% (F1-score), 7.0% (AUPRC), and 5.9% (AUROC). Overall, CLADES consistently improved predictive accuracy across both exon groups.

**Table 2:** Tabula Sapiens ELRC Performance. Performance comparison on the Tabula Sapiens dataset in the Exon-Level Regulation Classification (ELRC) task between CLADES and Baseline (no-CL) models for high- and low-expression exon groups. Metrics are reported per group using the 300 bp intron+exon configuration.

| Model | Exon Group | Precision | Recall | F1-score | AUPRC | AUROC |
| --- | --- | --- | --- | --- | --- | --- |
| CLADES | High | **0.178** | 0.586 | **0.228** | **0.365** | **0.627** |
| Baseline | High | 0.160 | **0.602** | 0.215 | 0.347 | 0.601 |
| CLADES | Low | **0.257** | **0.597** | **0.313** | **0.444** | **0.604** |
| Baseline | Low | 0.242 | 0.561 | 0.294 | 0.415 | 0.570 |

Together, these results demonstrate that CLADES captures cell-type-specific splicing variation and maintains robust predictive performance across a diverse set of cellular identities.

## 4 Discussion

In this work, we introduced CLADES, a contrastive learning framework that leverages evolutionary conservation as a principled source of augmentation for learning sequence-based representations of alternative splicing regulation. By treating orthologous exon–intron junctions as positive pairs, the model learns to unite embeddings of exons that share regulatory function while separating unrelated ones, yielding a representation space that captures conserved motif- and position-dependent signals. Our results demonstrate that these embeddings generalize effectively to downstream prediction of tissue- and cell-type–specific differential inclusion, improving *Δψ* correlation, direction-of-change classification, and robustness in low-sample settings relative to existing state-of-the-art models. These findings suggest that evolution encodes regulatory invariants that can be extracted via contrastive objectives, and that *cis*-regulatory programs governing context-dependent splicing possess a conserved structure recoverable from sequence alone. In addition to the empirical gains facilitated by orthologous positive pairs, we introduce two complementary problem formulations motivated by the biology of splicing regulation. We recast differential splicing as a direction-of-change classification task, formalizing TSRC and ELRC. We thus provide a more interpretable framework for evaluating regulatory outcomes.

Nonetheless, several limitations remain: orthology does not universally imply regulatory equivalence, especially for lineage-specific or rapidly evolving exons; fixed intronic windows may omit distal regulatory elements; and single-cell splicing measurements introduce substantial noise. Moreover, our current contrastive scheme uses simple uniform negative sampling and does not yet incorporate multimodal, multilevel regulatory information such as nucleosome position or RBP binding data. Future directions include phylogeny-aware augmentation strategies, integration of additional regulatory modalities, improved sampling schemes, and scaling CLADES toward a splicing regulatory foundation model capable of predicting splicing across conditions, perturbations, and evolutionary distances. Extending the framework to explicitly capture clade-specific, non-conserved regulatory programs represents an important direction for future investigation. Together, these results highlight the promise of contrastive, evolution-guided representation learning as a potentially general strategy for understanding biology using AI.

## Code Availability

The source code is available at https://github.com/ArghamitraT/CLADES.

## 5 Acknowledgements

We thank Karin Isaev and Alan Moses for their contributions. We acknowledge Empire AI Consortium (4) computing resources supported by Empire State Development, the Simons Foundation, and the Secunda Family Foundation. This work was supported by NSF CAREER DBI2146398, NSF CSGrad4US, and NIH Training Grant 5T15LM007079-34. The views expressed are those of the authors and do not necessarily reflect those of the NSF or NIH.

