## Supplementary Appendix for "CLADES — Contrastive Learning Augmented DifferEntial Splicing with Orthologous Positive Pairs"

### 1 Appendix

Table A1: **Summary of notations.**

| **Symbol** | **Definition** |
|---|---|
| $\psi_{e,t}$ | Percent spliced-in (PSI) of exon $e$ in tissue or cell type $t$ |
| $\overline{\psi}_e$ | Mean inclusion level of exon $e$ across tissues or cell types |
| $\Delta\psi_{e,t} = \psi_{e,t} - \overline{\psi}_e$ | Context-dependent change in exon inclusion |
| $\Delta\text{logit}(\psi_{e,t})$ | Change in inclusion on logit (log-odds) scale |
| $N, M$ | Anchors per batch and positives per anchor |
| $e_i$ | $i$-th anchor exon in the minibatch ($i = 1, \dots, N$) |
| $x_{i,m}^{(u)}$ | Input sequence of exon $i$, OPP $m$ at boundary $u \in \{5', 3'\}$ |
| $l_{\text{in}}, l_{\text{ex}}$ | Lengths of intronic and exonic regions used as input (bp) |
| $l_{\text{total}} = l_{\text{in}} + l_{\text{ex}}$ | Total boundary sequence length |
| $P(i)$ | Set of $M$ orthologous positives for anchor $e_i$ |
| $C(i)$ | All contrastive samples of anchor $i$ |
| $f_\theta^{(u)}$ | Encoder network for boundary $u$ |
| $y_{i,m}$ | Averaged representation and output of encoder, $\mathbb{R}^{d_{\text{feat}}}$ |
| $d_{\text{feat}}$ | Feature dimension (64) |
| $g_\phi$ | Projection head: $\mathbb{R}^{d_{\text{feat}}} \to \mathbb{R}^{d_{\text{proj}}} \to \mathbb{R}^{d_{\text{emb}}}$ |
| $d_{\text{proj}}$ | Projection layer dimension (512) |
| $d_{\text{emb}}$ | Final embedding dimension (128) |
| $z_{i,m}$ | Final projected embedding of exon $x_{i,m}^{(u)}$ |
| $\mathcal{L}_{\text{sup}}$ | Supervised contrastive loss |
| $\tau$ | Temperature hyperparameter |
| $\mathcal{L}_{\text{kl}}$ | Finetuning loss |
| $\rho$ | Spearman Correlation |
| $\Delta\psi^+$ | Up-regulated exons |
| $\Delta\psi^-$ | Down-regulated exons |
| $\Delta\psi_\emptyset$ | Unchanged exons |
| $\sigma_e$ | Standard deviation of exon e |

#### 1.1 Robustness to Incomplete Cross-Species Observations

**Robust Representation Learning under Noisy and Incomplete Cross-Species Observations.** Orthologous exons in CLADES are defined via genomic alignment and splice-site conservation. Although sequence alignment provides a biologically grounded notion of correspondence, it does not ensure that an exon is actively spliced or expressed in every species. This limitation primarily reflects incomplete RNA-seq and isoform-level annotations across non-model organisms and therefore represents a constraint of available cross-species data rather than a methodological shortcoming. Despite incomplete functional observations, the framework is designed to learn stable regulatory representations. This behavior is supported by Figure 3(a) and (b), where zero-shot embeddings from the contrastively pretrained encoder, without downstream fine-tuning, organize exons according to context-specific regulatory properties consistent with downstream predictive performance.

This constraint directly motivates the use of contrastive learning. Contrastive objectives are effective when views preserve task-relevant structure while attenuating nuisance variation. The InfoMin principle formalizes this by showing that representations emerge when views share information about the downstream signal but discard view-specific noise (9; 4). In CLADES, views are anchored at human exon–intron splice sites, which constitute one of the most evolutionarily conserved regulatory architectures across vertebrates (1; 6). Orthologous exon–intron contexts from other species serve as views in which conserved regulatory features form the shared signal reinforced during training, while species-specific inactivity or annotation noise remains view-specific and is attenuated.

CLADES further employs supervised contrastive learning, which improves robustness to distributional shift and input corruption relative to cross-entropy objectives (7). Aggregating multiple views per anchor reduces sensitivity to noisy positives and stabilizes alignment around shared regulatory structure. This effect is empirically reflected in the consistent performance gains observed as the number of views increases (Table 1). Prior theoretical and empirical analyses similarly demonstrate that contrastive learning promotes robustness to noisy labels by emphasizing consistent structure while suppressing noise-induced correlations (5; 10; 2). Thus, while alignment does not guarantee functional activity in every species, it provides a scalable and biologically principled notion of correspondence in settings lacking direct functional supervision.

**Correspondence Assumptions and Conservation Bias.** The CLADES framework is designed to limit reliance on curated annotations while avoiding conservation-induced bias. Curated resources are incorporated only at two narrowly scoped levels: reference definition and cross-species mapping. Genomic coordinates are defined exclusively using the human gene model. Human exon–intron boundaries represent among the most rigorously curated genomic features available (3) and are used solely to define training loci rather than as supervision targets. The model is never trained to predict conservation or annotation status, preventing preferential weighting of well-annotated events. Cross-species correspondence is established through multiple sequence alignment anchored at splice sites. Canonical splice sites are among the most evolutionarily constrained genomic elements due to mechanistic requirements of the splicing machinery (1; 6). This defines correspondence without assuming conservation of exon usage or regulatory strength. When regulatory mechanisms are lineage-specific or recently evolved, meaningful cross-species alignment is not expected (8). In such cases, the absence of alignment reflects biological divergence rather than methodological bias, and these events remain analyzable in downstream human-only tasks.

#### 1.2 ASCOT Dataset Additional Evaluation

(a)

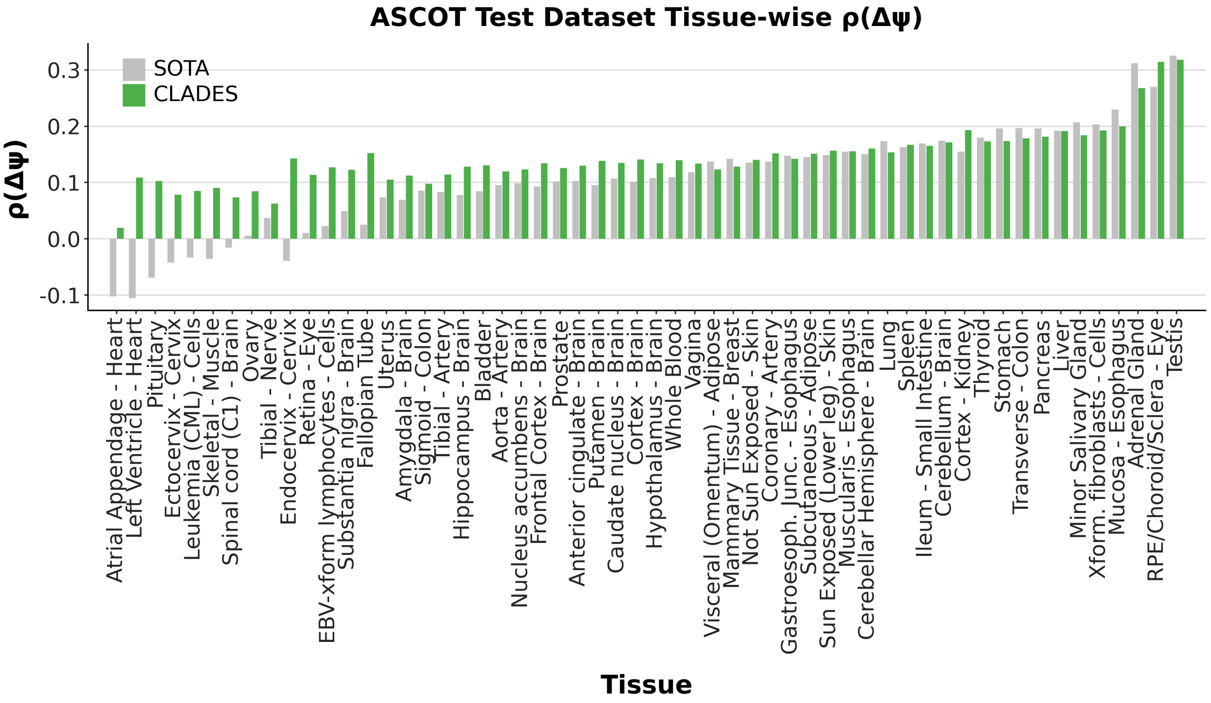


(b)

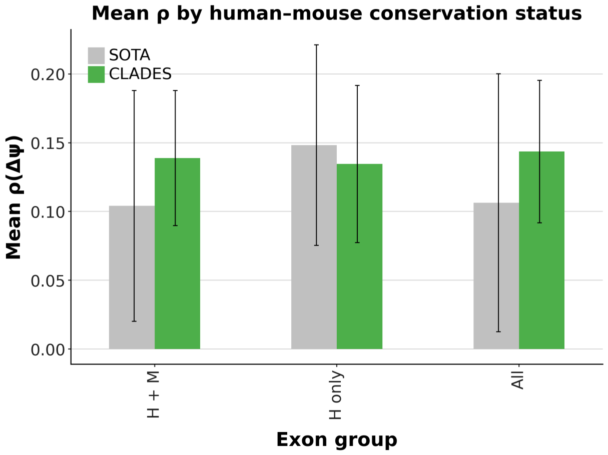


Fig. A1: **Evaluation on the held-out ASCOT test set.** Although we evaluate CLADES on the ASCOT variable-exon benchmark, we additionally follow the MTSplice data split and use the remaining 11,840 exons from chromosomes 2, 3, and 5 as an independent held-out test set. **(a)** Tissue-wise Spearman correlation $\rho(\Delta\psi)$ between the predicted and ground-truth tissue-dependent inclusion changes across all ASCOT tissues. Each bar corresponds to one tissue, with the baseline model shown in gray and CLADES shown in green; CLADES consistently improves correlation across the majority of tissues. **(b)** Exons in the held-out ASCOT test set were grouped according to human–mouse alignment status: 9,106 exons with detectable human–mouse alignment (H + M), 2,734 exons lacking detectable mouse sequence (H only), and All, which includes the full set of exons. Bars show mean $\rho(\Delta\psi)$ across 56 tissues, with error bars indicating $\pm 1$ standard deviation. The contrastively pretrained model outperforms the supervised baseline for exons with human–mouse alignment (H+M) and across the full test set (All), while remaining competitive but not superior for exons without detectable mouse sequence, likely due to weaker cross-species conservation signals.

Table A2: **Ablation with Embedding Dimensionality.** Performance comparison across CNN depth (2/4/6/8 layers) with and without contrastive pretraining on two evaluation datasets. The CLADES model corresponds to the 8-layer encoder configuration. Cells report Spearman correlation $\rho(\Delta\psi)$.

| **Dataset** | **Training** | **CNN-2** | **CNN-4** | **CNN-6** | **CNN-8** |
|---|---|---|---|---|---|
| Variable Exon Set | No CL | 0.254 | 0.237 | 0.238 | 0.226 |
| | W/ CL | 0.276 | 0.276 | 0.276 | **0.284** |
| Test Exon Set | No CL | 0.107 | 0.104 | 0.099 | 0.106 |
| | W/ CL | 0.147 | **0.151** | 0.149 | 0.144 |

We include an ablation study in Table A2 evaluating the effect of encoder depth on performance, reporting mean Spearman correlation $\rho(\Delta\psi)$ the variable and the test exon set. While the main results in the paper use an encoder with eight CNN blocks, we also evaluate models with two, four, and six blocks. Each CNN block consists of a convolutional layer followed by a residual connection and batch normalization. As shown in Table A2, contrastive pretraining consistently improves performance across all encoder depths and both evaluation datasets. Notably, the strongest performance on the variable exon set is achieved by the CNN-8 model trained with contrastive learning ($\rho = 0.284$), which corresponds to the primary CLADES architecture used throughout the paper. A similar trend is observed on the test exon set, where contrastive models consistently achieve higher $\rho$ than non-contrastive baselines.

##### 1.3 Cell-Specific Tabula Sapiens Dataset:

**(a)**

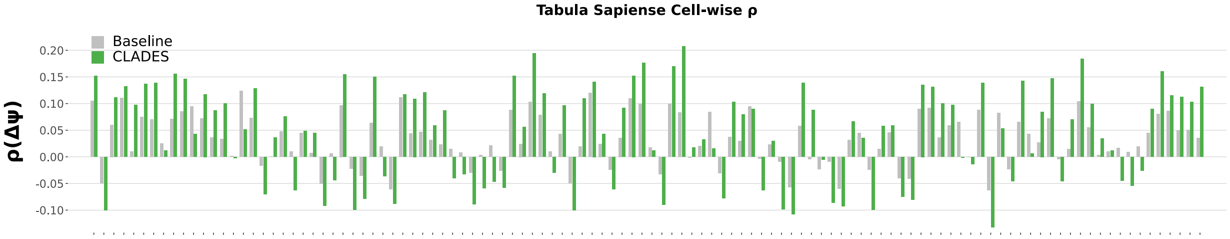


**(b)**

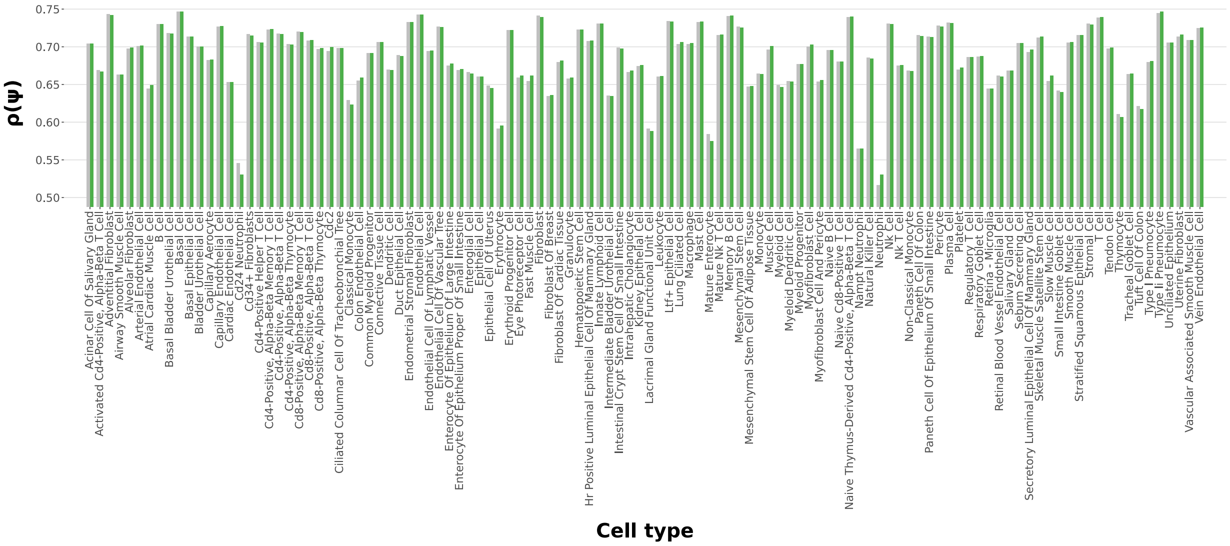


Fig. A2: **Cell-specific evaluation on the Tabula Sapiens dataset.** To assess the generalization of CLADES beyond bulk-tissue splicing patterns, we evaluated the model on the Tabula Sapiens single-cell atlas. For each cell type, we computed Spearman correlation ($\rho$) between predicted and observed $\Delta\psi$ and $\psi$. **(a)** Cell-type–wise Spearman correlation between predicted and observed differential inclusion $\rho(\Delta\psi)$. Each bar corresponds to one annotated Tabula Sapiens cell type, with the baseline model shown in gray and CLADES shown in green. Performance varies substantially across cell types, yet CLADES improves correlation in the majority of them. **(b)** Cell-type–wise Spearman correlation $\rho(\psi)$ for predicting absolute inclusion rather than differential changes. Here CLADES performs comparably to the baseline across most cell types, with modest improvements in several epithelial and immune populations.

**(a)**

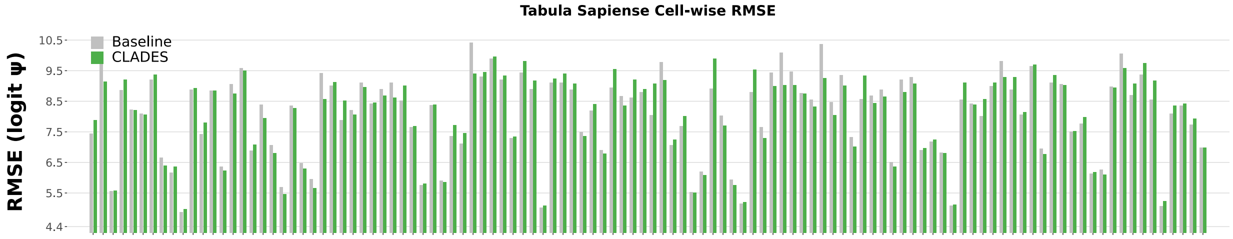


**(b)**

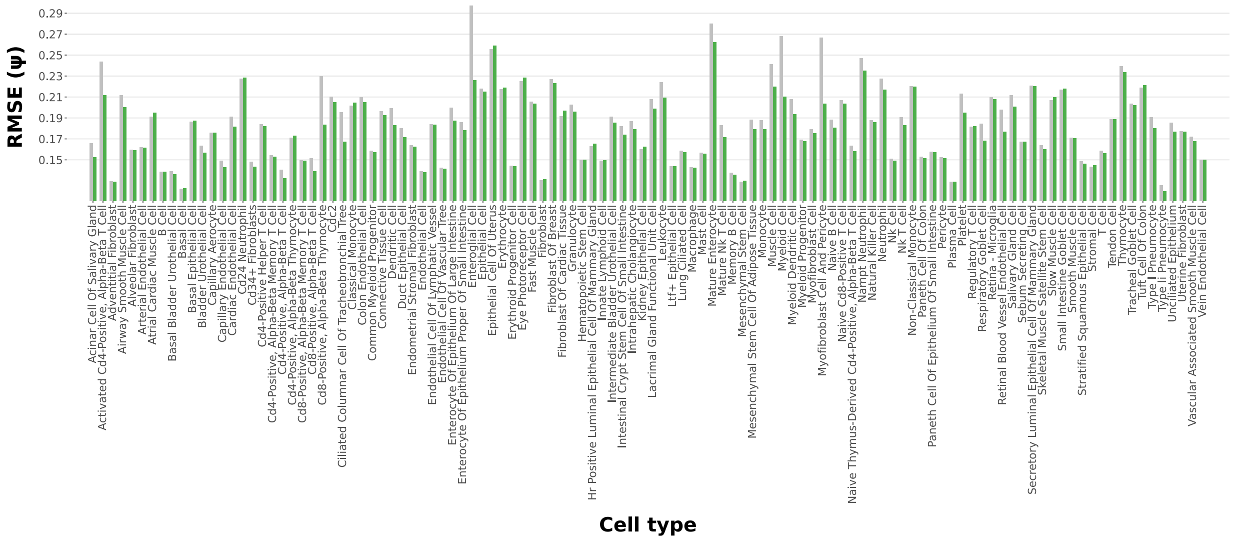


Fig. A3: **Cell-specific RMSE evaluation on the Tabula Sapiens dataset.** To evaluate how well CLADES generalizes to single-cell splicing variation, we assessed prediction error across all annotated Tabula Sapiens cell types. For each cell type, we computed root-mean-squared error (RMSE) between predicted and observed $\psi$, using either logit-transformed PSI or raw PSI depending on the task. **(a)** Cell-type–wise RMSE for predicting logit-transformed PSI. Each bar corresponds to one cell type, with the baseline model shown in gray and CLADES in green. CLADES achieves consistently lower error across a wide range of epithelial, immune, and stromal populations. **(b)** Cell-type–wise RMSE for predicting raw PSI. Similar trends are observed: CLADES generally reduces prediction error relative to the baseline, with notable improvements in several highly variable immune and endothelial populations.

##### 1.4 Dataset Statistics

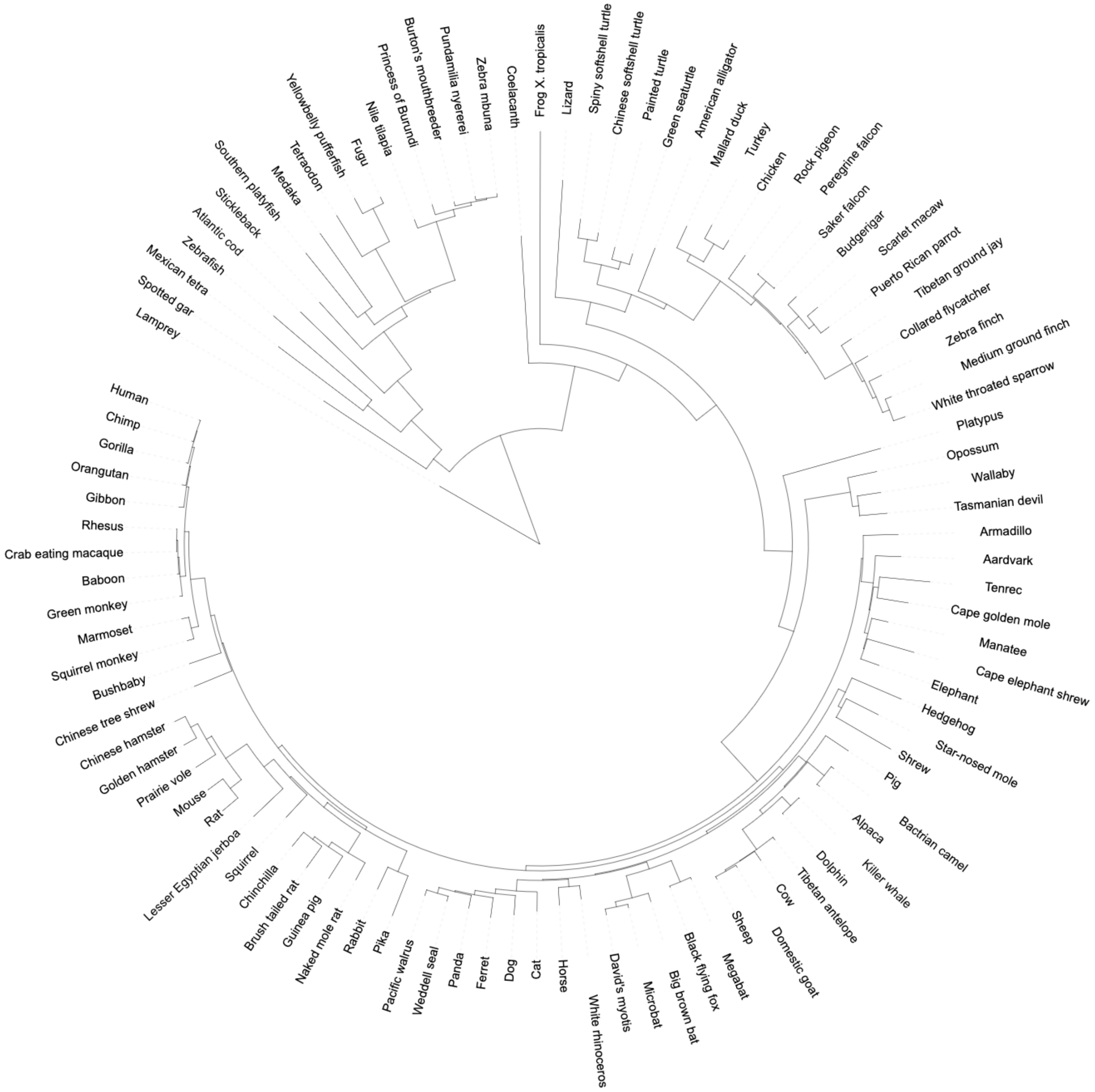


Fig. A4: **Multiz100way phylogenetic tree.** The pretraining dataset leverages the Multiz100way MSA, and the figure shows the phylogenetic relationships among the species included in this alignment.

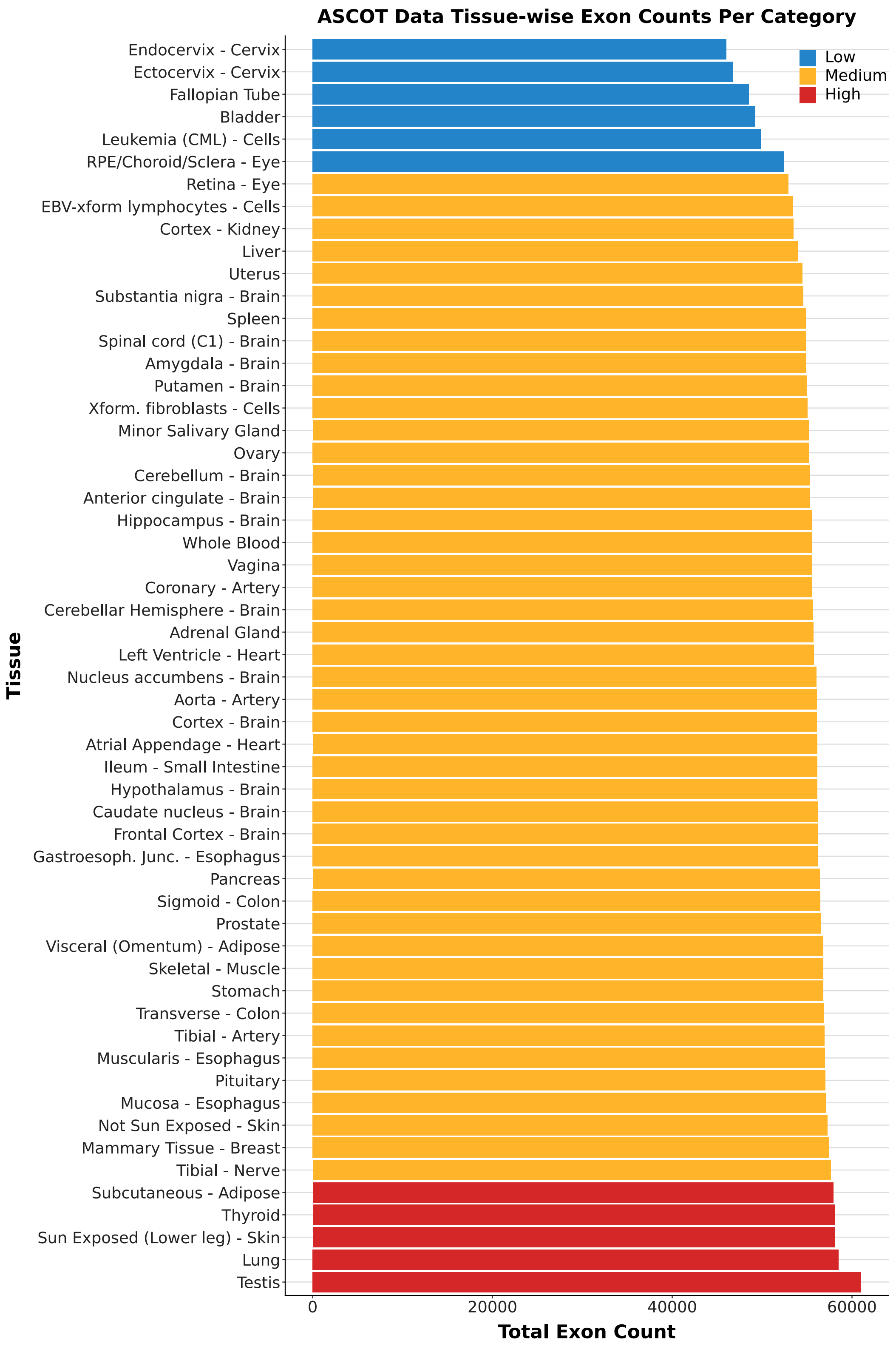


Fig. A5: **Tissue-wise exon counts in the ASCOT dataset.** Barplot showing the total number of exons in each tissue, stratified into low, medium, and high count categories.

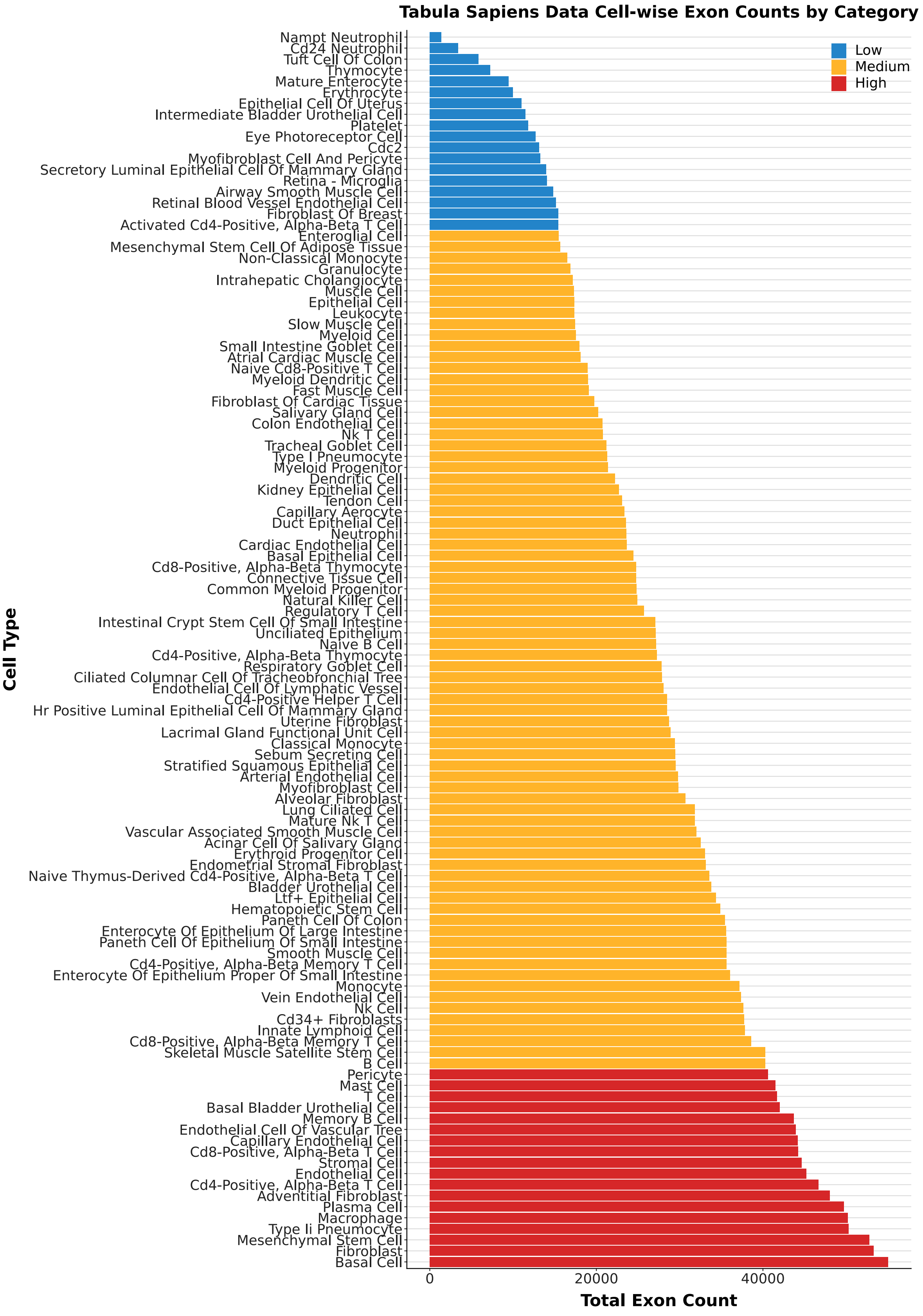


Fig. A6: **Cell-type–wise exon counts in the Tabula Sapiens dataset.** Barplot showing total exon counts per cell type, stratified by low, medium, and high observation categories.

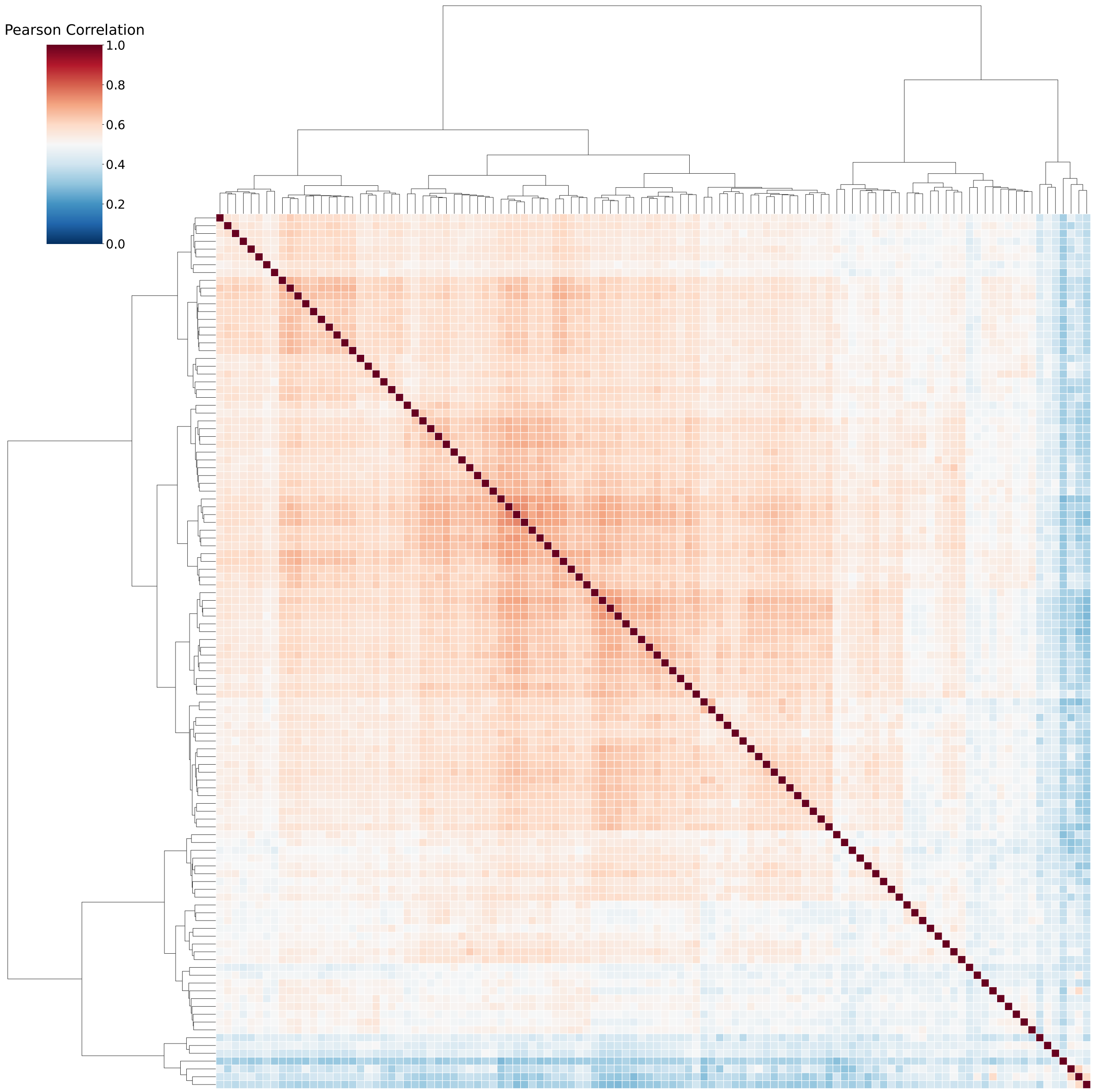


Fig. A7: **Tabula Sapiens cell–cell correlation structure.** The Tabula Sapiens dataset was filtered to include only exons that had at least 10 tissues where $\psi_{e,t}$ differed from $\overline{\psi}_e$ by 10% points. Then, the Pearson correlation between cell types was calculated and shown in this heatmap, where high correlation is shown in red and low correlation is shown in blue. Finally, Ward linkage was calculated for hierarchical clustering. Labels for each cell type are listed below ( 1.4)

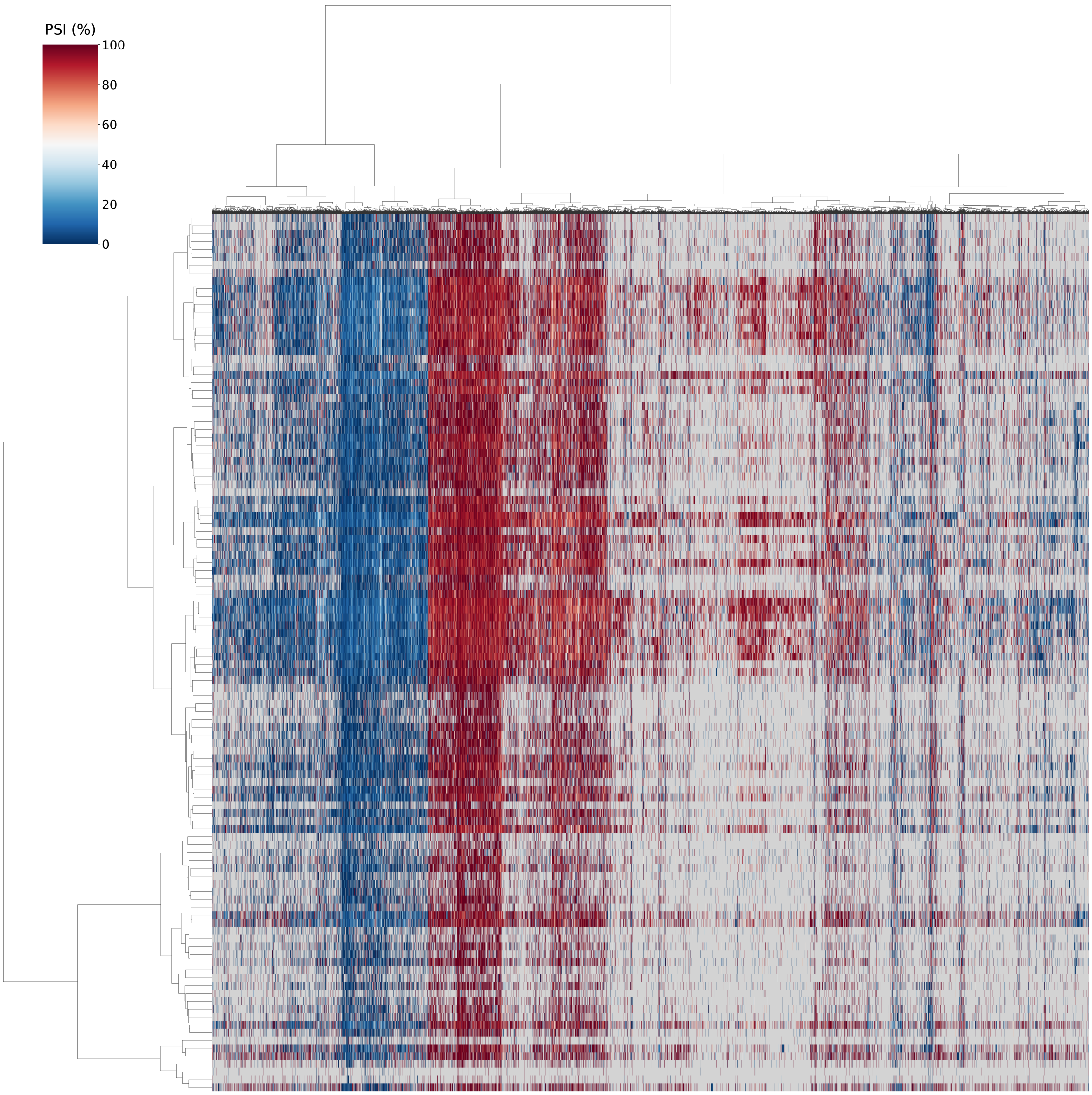


Fig. A8: **Tabula Sapiens exon–cell type heatmap.** A heatmap of $\psi_{e,t}$ for the 20,145 exons (columns) across each cell type (rows). Both exons and cell types were clustered using Ward linkage, where missing exons were imputed with $\overline{\psi}_e$. See previous figure for cell type clustering information. $\psi_{e,t}$ values are color coded from 0% (blue) to 100% (red). Gray values represent missing exons which rMATS did not detect. Labels for each cell type are listed below ( 1.4)

##### Cell Type Cluster Ordering

1. NK T Cell
2. Naive CD8-Positive T Cell
3. CD4-Positive, Alpha-Beta Thymocyte
4. CD8-Positive, Alpha-Beta Thymocyte
5. Natural Killer Cell
6. CD4-Positive Helper T Cell
7. Activated CD4-Positive, Alpha-Beta T Cell
8. Naive B Cell
9. NK Cell
10. CD4-Positive, Alpha-Beta T Cell
11. Memory B Cell
12. Innate Lymphoid Cell
13. B Cell
14. T Cell

15. CD8-Positive, Alpha-Beta Memory T Cell
16. CD8-Positive, Alpha-Beta T Cell
17. Naive Thymus-Derived CD4-Positive, Alpha-Beta T Cell
18. CD4-Positive, Alpha-Beta Memory T Cell
19. Myeloid Dendritic Cell
20. Dendritic Cell
21. Macrophage
22. Mature NK T Cell
23. Mast Cell
24. Regulatory T Cell
25. Intestinal Crypt Stem Cell of Small Intestine
26. Ciliated Columnar Cell of Tracheobronchial Tree
27. Lung Ciliated Cell
28. Respiratory Goblet Cell
29. Paneth Cell of Epithelium of Small Intestine
30. Paneth Cell of Colon
31. Unciliated Epithelium
32. Bladder Urothelial Cell
33. HR Positive Luminal Epithelial Cell of Mammary Gland
34. Sebum Secreting Cell
35. Duct Epithelial Cell
36. Intrahepatic Cholangiocyte
37. LTF+ Epithelial Cell
38. Stratified Squamous Epithelial Cell
39. Basal Cell
40. Type II Pneumocyte
41. Kidney Epithelial Cell
42. Basal Bladder Urothelial Cell
43. Acinar Cell of Salivary Gland
44. Erythroid Progenitor Cell
45. Plasma Cell
46. Hematopoietic Stem Cell
47. Myeloid Progenitor
48. Common Myeloid Progenitor
49. CD34+ Fibroblasts
50. Fibroblast
51. Mesenchymal Stem Cell
52. Adventitial Fibroblast
53. Pericyte
54. Stromal Cell
55. Capillary Endothelial Cell
56. Endothelial Cell of Vascular Tree
57. Endothelial Cell
58. Arterial Endothelial Cell
59. Vein Endothelial Cell
60. Endothelial Cell of Lymphatic Vessel
61. Type I Pneumocyte
62. Mesenchymal Stem Cell of Adipose Tissue
63. Slow Muscle Cell
64. Fast Muscle Cell
65. Retinal Blood Vessel Endothelial Cell
66. Cardiac Endothelial Cell
67. Basal Epithelial Cell
68. Capillary Aerocyte
69. Fibroblast of Cardiac Tissue
70. Myofibroblast Cell
71. Endometrial Stromal Fibroblast
72. Uterine Fibroblast
73. Atrial Cardiac Muscle Cell
74. Vascular Associated Smooth Muscle Cell
75. Smooth Muscle Cell
76. Airway Smooth Muscle Cell
77. Alveolar Fibroblast
78. Tendon Cell
79. Skeletal Muscle Satellite Stem Cell
80. Platelet
81. Eye Photoreceptor Cell
82. Muscle Cell
83. Colon Endothelial Cell
84. Connective Tissue Cell
85. Myofibroblast Cell and Pericyte
86. Enteroglial Cell
87. Fibroblast of Breast
88. Salivary Gland Cell
89. Small Intestine Goblet Cell
90. Enterocyte of Epithelium of Large Intestine
91. Enterocyte of Epithelium Proper of Small Intestine
92. cDC2
93. Intermediate Bladder Urothelial Cell
94. Epithelial Cell
95. Secretory Luminal Epithelial Cell of Mammary Gland
96. Tracheal Goblet Cell
97. Epithelial Cell of Uterus
98. Erythrocyte
99. Leukocyte
100. Thymocyte
101. Retina - Microglia
102. Myeloid Cell
103. Non-Classical Monocyte
104. Monocyte
105. Granulocyte
106. Tuft Cell of Colon
107. Classical Monocyte
108. Lacrimal Gland Functional Unit Cell
109. Mature Enterocyte
110. NAMPT Neutrophil
111. CD24 Neutrophil
112. Neutrophil
